# Direct thin-layer agar for bedaquiline-susceptibility testing of *Mycobacterium tuberculosis* at BSL2 level yields high accuracy in 15 days from sputum processing

**DOI:** 10.1101/2024.11.18.624108

**Authors:** I. Cuella Martin, D. Runyambo, S. De Bock, M. Diels, H. Niyompano, F. Hakizayezu, J. Keysers, W.B. De Rijk, Y. M. Habimana, N. Gahamanyi, E. Ardizzoni, C.M. Muvunyi, B.C. de Jong, J.C.S. Ngabonziza, L. Rigouts

## Abstract

This study evaluated thin-layer agar (TLA) as a faster alternative for both indirect MIC determination of bedaquiline (BDQ) from culture isolates and direct drug-susceptibility testing (DST) from sputum samples. Indirect BDQ-MIC results from TLA were compared to the established 7H11 solid DST. Direct-TLA-DST performance was assessed using 143 baseline sputum samples from rifampicin-resistant TB cases. Direct-TLA tested susceptibility to rifampicin, isoniazid, levofloxacin, and BDQ, with results compared to Löwenstein-Jensen (LJ) and MGIT.

For indirect BDQ-MIC determination, TLA demonstrated high accuracy, with MIC values for H37Rv falling within the WHO control range (0.015-0.12μg/ml). The incubation conditions (5-10% CO_2_, standard) and reading time (day 7, 14) significantly affected results, with D7 readings in standard incubator providing optimal accuracy. Direct-TLA-DST detected MTB in 53.8% of samples, compared to 55.9% on LJ and 69.4% in MGIT. Uninterpretable results due to contamination or medium drying were low (4.9%). Median time to result was 15 days for smear-positive and 22 days for smear-negative samples, faster than WHO endorsed methods. Sensitivity was 100% for rifampicin and 87.8% for isoniazid, with a specificity of 100% for all drugs except isoniazid (96.2%). No BDQ nor levofloxacin resistance was detected, thus direct TLA sensitivity could not be assessed.

In conclusion, direct-TLA-DST offers a reliable and faster alternative to conventional DST methods for BDQ and other anti-TB drugs. Essentially, this technique can be operated at BioSafety Level 2, allowing decentralization of pDST for managing drug-resistant TB in settings with limited laboratory infrastructure.

**Importance:** This paper addresses the critical need for faster direct drug-susceptibility testing (DST) on sputum, especially for bedaquiline (BDQ), which is a key drug in treating drug-resistant TB. Currently, there is a lack of rapid, reliable methods for direct BDQ testing from sputum samples, limiting timely and accurate treatment decisions and monitoring. By demonstrating the potential of thin-layer agar (TLA) for direct BDQ-MIC determination, this study offers a promising solution that could significantly improve patient care.

## INTRODUCTION

Management of multidrug-resistant tuberculosis (MDR-TB) is compromised by limited access to drug-susceptibility testing (DST), which is increasingly crucial to optimise care for MDR-TB patients.^(1)^ Since 2020, the World Health Organization (WHO) has included bedaquiline (BDQ) as a core component of any MDR-TB regimen.^(2)^ BDQ is a diarylquinoline antimycobacterial compound that exerts its bactericidal effect by blocking ATP synthesis.^(3)^ Although BDQ is a relatively new drug, increasing rates of resistance are being reported, including among drug-naïve TB patients, which compromises these regimens.^(4,5)^

Only few patients who start an all-oral BDQ based regimen today will have resistance testing done for BDQ, as phenotypic testing requires a BSL3 laboratory, and genotypic testing has not yet been translated to a rapid molecular test, due to the imperfect correlation of a wide range of mutations with phenotypic resistance. Resistance to BDQ has been associated with genetic mutations or resistance-associated variants (RAVs) in the *atpE, Rv0678*, and *pepQ* genes.^(6,7)^ In vitro studies indicate that mutations in *atpE*, the gene encoding BDQ’s direct target ATP synthase, lead to high-level BDQ resistance, although these mutations likely cause significant fitness loss and hence are rarely found in clinical isolates.^(8–10)^ In contrast, mutations in the *Rv0678* efflux pump regulator gene are linked to low-level BDQ resistance, but only if the efflux pump remains functional.^(11–14)^ Mutations in *pepQ* generally result in a modest increase in BDQ minimum inhibitory concentration (MIC).^(13,15)^ Given these imperfect genotypic–phenotypic associations phenotypic DST (pDST) remains the reference test to effectively guide treatment and monitor the emergence of BDQ resistance.

The current guidelines for phenotypic BDQ-susceptibility testing are based on limited evidence.^(16)^ Although MIC determination using Middlebrook 7H10 or 7H11 agar and 7H9 broth microdilution method (BMD) have been validated in a multicountry study, solid media-based DST is restricted to polystyrene tubes and time-consuming, often requiring three to four weeks to yield results after a positive primary culture is obtained.^(17)^ Liquid-based methods, such as the Mycobacteria Growth Indicator Tube (MGIT) system, are recommended for their higher sensitivity and faster turnaround times.^(18)^ However, all these methods require BSL3 biosafety infrastructure and equipment, limiting decentralized access, particularly where specimen transport systems to centralized laboratories are inadequate.

Thin-layer agar (TLA) offers a simpler and more afordable alternative for isolating *Mycobacterium tuberculosis* (MTB) complex and detecting drug resistance directly from sputum samples. TLA has demonstrated comparable performance to conventional methods, such as the MGIT system, for MTB isolation and sensitivity and specificity for the detection of rifampicin (RIF) and isoniazid (INH) resistance.^(19,20)^ Its one-step approach and suitability for biosafety level 2 (BSL2) laboratories make it a promising option for use in district or peripheral settings.^(21)^ However, the use of TLA for BDQ testing has not been documented.

This study aimed (1) to validate the performance of the TLA as an indirect method for BDQ-MIC determination, comparing it with the standard tube-based 7H11 MIC method, and (2) to evaluate the performance of direct TLA-DST under various conditions for MTB isolation and pDST for BDQ, along with RIF, INH, and levofloxacin (LFX), in comparison to solid and liquid culture followed by indirect pDST.

## MATERIALS AND METHODS

### Indirect TLA-BDQ-MIC validation

Twenty-one clinical MTB isolates and 10 in-vitro-selected strains obtained from the collection at the Institute of Tropical Medicine (ITM) in Antwerp, Belgium, were used for validation of indirect TLA-MIC determination for BDQ (TLA-BDQ-MIC) (Table S1). Of the 21 clinical isolates, WGS data was available for 18, of which all had a wild-type *atpE* gene, while three had a mutation in the *Rv0678* gene. Four of the 10 in-vitro-selected strains presented a mutation in *atpE* and six in *Rv0687*. The MTB H37Rv reference strain (ITM 500735) was included as BDQ-susceptible control strain. All isolates were freshly subcultured on Löwenstein-Jensen (LJ) medium before being tested.

A bacterial suspension prepared from fresh MTB growth on LJ was adjusted to a turbidity of McFarland standard No. 1 and further diluted 1:10 in sterile distilled water. Using a sterile transfer pipette, one drop (∼50µl) was inoculated into each of the 12 wells of an in-house prepared TLA plate (Corning Costar 3513; Sigma-Aldrich, USA), except for the negative control (NC) well. Two operators independently performed the inoculation in duplicate, starting from a different bacterial suspension, for a total of 4 replicates. Each plate consisted of one well for growth control (GC), one well for p-nitrobenzoic acid (PNB) at 500µg/ml to differentiate between MTB complex and non-tuberculous mycobacteria (NTM), one well for the NC, and one well for each serial concentration of BDQ: 0.008, 0.015, 0.03, 0.06, 0.125, 0.25, 0.5, 1 and 2 µg/ml. BDQ powder was obtained from Janssen Pharmaceuticals, Belgium, batch 16175328. To prepare the plates, Middlebrook 7H11 agar base (212203, Becton Dickinson, USA) was mixed with glycerol 86-89% and sterile distilled water as per the manufacturer’s instruction. After autoclaving and cooling down to 55°C the medium was supplemented with oleic acid dextrose catalase (OADC, 212240, Becton Dickinson) plus the antibiotics amphotericin, piperacillin, and trimethoprim (Sigma-Aldrich), all at 4 µg/ml concentration. Parallel MIC determination on 7H11 agar medium enriched with OAD supplement in polystyrene tubes was performed, using the same BDQ concentrations.^(18)^ After inoculation, parallel plates were placed in a 34-38°C incubator with ambient CO_2_ concentrations or supplemented with 5-10% CO_2_, while tubes were always incubated with 5-10% CO_2_. TLA plates were read on days 7 (D7) and 14 (D14) by two independent operators using a conventional light microscope (objective 10x). Growth in the GC well paired with the absence of growth in the PNB well was considered positive for MTB. The MIC was interpreted if at least 10 colonies were present in the GC. MIC results corresponded to the well with the lowest drug concentration that did not show growth, i.e. 100% inhibition. For tube-based 7H11 MIC testing, reading was done at 4 weeks, and the MIC_99_ was determined as the highest concentration that had less growth than the 1/100 diluted GC tube.

### Direct on sputum TLA-DST

We collected sptum samples from consecutive patients diagnosed with pulmonary rifampicin-resistant (RR-) TB by GeneXpert MTB/RIF (Ultra) who were registered in the MDR-TB Clinic in Kabutare between December 2021 and December 2023 and who consented to participate in the study. Three sputum samples were collected before starting RR-TB treatment: the first on the day of arrival (spot sample 1), the second collected overnight (sample 2; sputum accumulated in the same container throughout the night), and the third on the following day (spot sample 3). All three samples were sent together on the day the third sample was obtained, using a cool box, to the National Reference Laboratory (NRL) in Kigali, Rwanda, for further genotypic and phenotypic testing. After NaOH-Nalc decontamination, the three samples underwent Auramine-O microscopy. Additionally, sample two was used for primary culture on homemade LJ and in the automated MGIT, and sample three for a second primary culture (LJ and MGIT) and parallel direct-TLA-DST. Leftovers of the three sediments were stored at -20°C. Subsequent pDST for positive LJ or MGIT cultures was done either by the 1% proportion method on LJ medium against first-line TB drugs or, since October 2022, by MGIT-DST including first-line drugs, BDQ, and levofloxacin (LFX). Direct-TLA testing was performed using the same 12-well plates (brand, type) and 7H11 agar medium supplemented with a broad-spectrum antibiotic mixture (i.e. OADC, amphotericin, piperacillin, and trimethoprim) to suppress contamination, mirroring the procedures previously detailed at ITM. In addition, some plates were prepared with the addition of the redox indicator 2,3-diphenyl-5-(2-thienyl) tetrazolium chloride (STC; Merck, USA) at a final concentration of 50 µg/ml, which was incorporated into the medium after cooling down to potentially facilitate plate reading and interpretation of observed (early) growth.^(22)^ To ensure the quality and consistency of each batch of in-house prepared plates, a quality control check was performed by inoculating H37Rv and *Mycobacterium fortuitum*. Each 12-well plate could accomodate two samples with for each sample a drug-free GC and wells containing each PNB (500 µg /ml), RIF (1.0 µg /ml), INH (0.2 µg /ml), LFX (1.0 µg/ml) or BDQ (0.25 µg/ml). The processed sputum sediments were resuspended in 2 mL of phosphate-buffered saline (PBS). A sterile transfer pipette was used to inoculate approximately 50 µL (one drop) of the resuspended sample into each TLA well. Plates were incubated at 34–38°C in a CO_2_-enriched environment, generated using a candle jar, and visually examined without opening on days 7, 14, 21, 28, and 35, or the nearest available day if deviations from the schedule occurred. From May 2023 onwards, TLA plates were incubated under standard incubation conditions without CO2 supplementation. Results were interpreted on the day of GC positivity. A positive result for MTB was defined by the absence of growth in the PNB well, while drug resistance was indicated by any growth in drug-containing wells. Discrepancies across all drug results between direct-TLA-DST and indirect DST performed on LJ or MGIT were resolved through gene sequencing. Sanger sequencing of the *rpoB* gene was conducted on residual sputum or sediment samples (second or third sample) that were shipped to the Institute of Tropical Medicine (Antwerp, Belgium).^(23)^ Additionally, sequencing of the *katG* and *fabG-inhA* genes was performed to assess isoniazid (INH) resistance. Whole-genome sequencing (WGS) was performed on genomic DNA extracted from isolates using the Illumina HiSeq platform (CD Genomics, USA) to further resolve discrepancies for INH, LVX, and BDQ(6). In cases where sequencing data were unavailable to resolve discrepancies, the results from indirect pDST were considered the definitive result.

### Statistical analysis

A Chi-square test with Yates’ continuity correction was conducted to compare the accuracy of TLA-BDQ-MIC results across different incubation environments and reading times. Additionally, a Z-test for proportions was employed to evaluate the significance of differences in result reliability when comparing standard incubator readings at day 7 to other conditions. The MTB positivity rate for direct TLA-DST was determined by dividing the number of samples exhibiting MTB by the total samples tested. Different conditions for direct TLA method were compared by Fisher’s exact test using LJ results as a baseline through Mantel-Haenszel chi-squared test. TLA results were compared with those from LJ or MGIT using matched sample IDs and the Z proportion method, which was adjusted for multiple comparisons with a Bonferroni post-hoc correction. Sensitivity and specificity, along with their corresponding 95% confidence intervals (CI), were calculated for TLA in detecting resistance to RIF, INH, LFX, and BDQ, using LJ and MGIT as reference standards, based on availability. All statistical analyses were conducted using R software, version 4.1.0, with a significance level set at 0.05.

### Ethics approval

The study protocol was approved by the Institutional Review Board of ITM (IRB/AB/AC/157; Ref No.1525/21; 23/09/2021), the Ethics Committee of the University Hospital of Antwerp, Belgium (REG No. B3002021000230; 22/11/2021), and the Rwanda National Ethics Committee (IRB 00001497 of IORG0001100; Ref No·705/RNEC/2021).

## RESULTS

### Indirect TLA-BDQ-MIC validation

For the H37Rv strain, BDQ-MIC values were slightly elevated under 5-10% CO_2_ incubation with a mode of 0.125 ug/ml and a range of 0.060-0.125 ug/ml, compared to a mode of 0.060 ug/ml and a range of 0.030 - 0.125 ug/ml under standard incubation conditions (Figure 1). In both incubation environments, a trend towards higher MIC values was observed at D14 reading compared to D7. Notably, among all H37Rv replicates, only one plate exhibited contamination at day 7 in the standard incubator, accounting for 0.9% (1/112).

**Figure 1.**
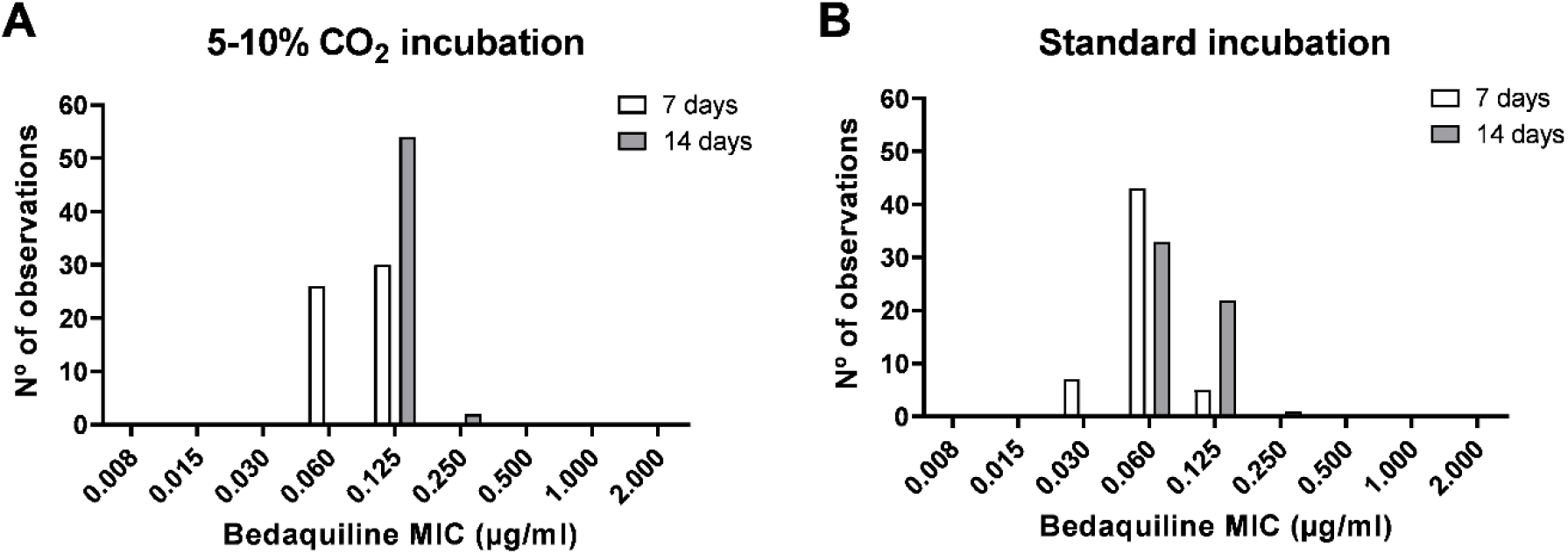
Minimal inhibitory concentration (MIC) distribution of bedaquiline for 56 replicates of the *M.tuberculosis* H37Rv reference strain on thin layer agar plates incubated in parallel at various conditions, with reading after 7 days and 14 days.

**Figure 2.**
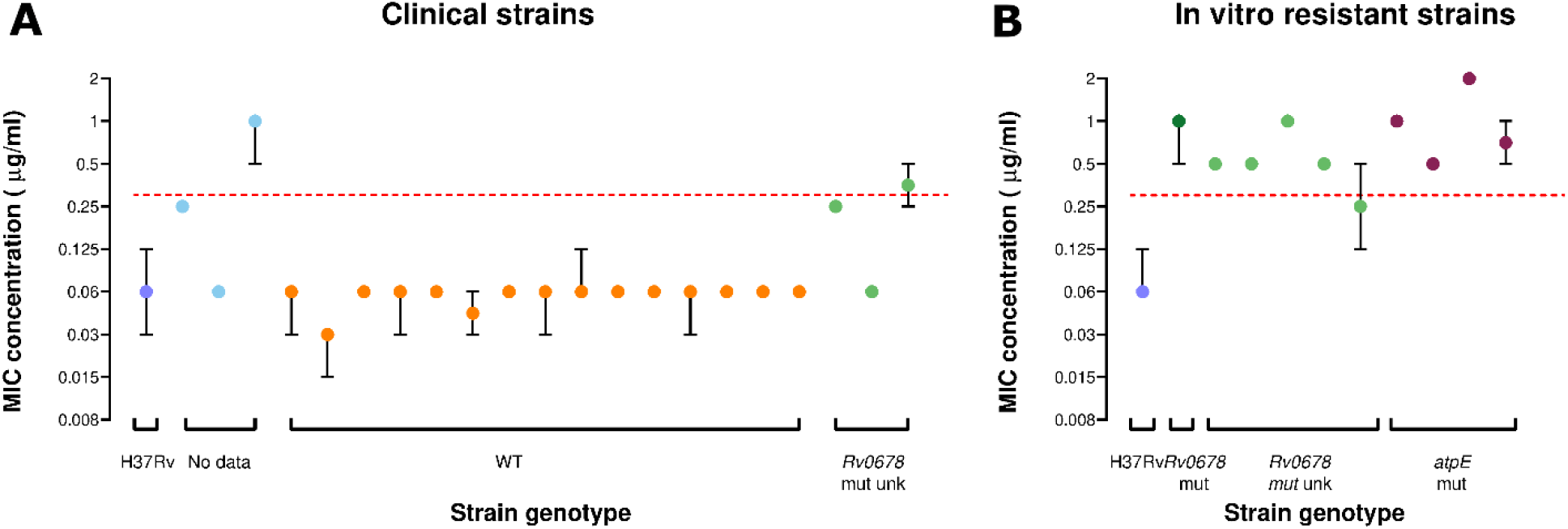
Bedaquiline (BDQ) minimal inhibitory concentration (MIC) median distribution of replicates with 95% CI of (A) 21 clinical *M. tuberculosis* isolates and (B) 10 in-vitro resistant strain, tested by indirect thin layer agar with standard incubation and reading at 7 days, stratified by genetic profile coded by color. H37Rv is included as control. The WHO proposed critical concentration for BDQ on 7H11 agar is marked with a dashed line at 0.25 µg/ml. *WT: wild type. Rv0678 mut unk: mutation in the Rv0678 gene that is either not associated with BDQ resistance or not listed in the second edition of the WHO mutation catalogue. Rv0678 mut: mutation in the Rv0678 gene that is associated with BDQ resistance. atpE mut: mutation in the atpE gene that is associated with BDQ resistance*.

For the 21 clinical isolates and 10 in vitro-resistant strains analyzed, TLA-BDQ-MIC determination was influenced by both the incubation environment and the reading time (Table 1). Results obtained from the standard incubator demonstrated significantly greater accuracy compared to the reference 7H11-tube testing than those from the 5–10% CO_2_ incubator, both at D7 (χ^2^ = 4.54, p = 0.033) and D14 (χ^2^ = 20.57, p < 0.001). The findings indicate that utilizing the standard incubator and conducting readings on day 7 improved the reliability of TLA-BDQ-MIC results, making this combination the preferred approach for optimal accuracy (Z = 2.38, p = 0.017) (Table 2). Of the 248 TLA plates, only 1 (0.40%) plate at D7 and 3 (1.21%) plates at D14 exhibited partial contamination, preventing the interpretation of the MIC results.

**Table 1.** Indirect TLA-BDQ-MIC results relative to 7H11 tube-based MIC testing, for 21 clinical isolates and 10 in-vitro resistant strains, with for 4 replicates per isolate/strain, stratified per incubation environment and day 7 (D7) or day 14 (D14) reading. Total number of plates tested equals 124. *MIC: minimum inhibitory concentration*.

| Number (%) of plates with MIC result relative to the 7H11 tube-based testing |  |  |  |  |  |  |
| --- | --- | --- | --- | --- | --- | --- |
| Reading time | 5-10% CO<sub>2</sub> incubator |  |  | Standard incubator |  |  |
|  | ≤ 1-fold dilution | > 1-fold dilution | Not interpretable | ≤ 1-fold dilution | > 1-fold dilution | Not interpretable |
| D7 | 112 (90.3%) | 11 (8.9%) | 1 (0.8%) | 121 (97.6%) | 3 (2.4%) | - |
| D14 | 91 (73.4%) | 31 (25%) | 2 (1.6%) | 118 (95.2%) | 5 (4%) | 1 (0.8%) |

**Table 2.** Overview of 16 samples with discordant drug-susceptibility testing (DST) results between direct thin-layer agar (TLA) and indirect DST on Löwenstein-Jensen or in MGIT medium, with sequencing results for relevant genes. In case no sequencing results were available, the indirect-DST results were considered as final results. *RIF: rifampicin. INH: isoniazid. LFX: levofloxacin R: resisitant. S: susceptible. WT: wild type. NT: not resolved*.

| Sample ID | Drug tested | Indirect DST | TLA | Sequencing | Final Result | In favor of direct-TLA testing |
| --- | --- | --- | --- | --- | --- | --- |
| 183 | RIF | R | S | *rpoB*_WT | S | Yes |
| 195 | RIF | S | R | *rpoB*_Asp435Ty | R | Yes |
| 201 | RIF | S | R | *rpoB*_Asp435Ty | R | Yes |
| 208 | RIF | R | S | *rpoB*_WT | S | Yes |
| 018 | INH | S | R | *katG*_Ser315Thr | R | Yes |
| 024 | INH | S | R | *katG*_Ser315Thr | R | Yes |
| 100 | INH | R | S | *katG*_Ser315Thr | R | No |
| 143 | INH | S | R | *katG*_Ser315Thr | R | Yes |
| 147 | INH | R | S | *fab1G*_15C>T | R | No |
| 183 | INH | R | S | WT | S | Yes |
| 200 | INH | R | S | *fab1G*_15C>T | R | No |
| 201 | INH | S | R | WT | S | No |
| 208 | INH | R | S | **NT** | R | **No** |
| 210 | INH | R | S | **NT** | R | **No** |
| 143 | LFX | R | S | WT | S | Yes |
| 222 | LFX | R | S | **NT** | R | **No** |

### Direct TLA-DST

From December 2021 to December 2023, a total of 143 sputum samples were received at the NRL in Rwanda from patients nationwide with a positive RR-TB result from the GeneXpert test conducted at local health facilities (Figure 3). The overall culture positivity rates prior to treatment start were 53.8% (77/143) for TLA, 55.9% (80/143) for LJ, and 69.4% (75/108) for MGIT. Among the 58 smear-microscopy-positive (sm+) samples, TLA’s performance in isolating MTB was comparable to LJ (82.8% vs. 87.9%; p=0.43) and MGIT (82.8% vs. 93.6%; p=0.093). For 85 smear-negative (sm-) samples TLA performed equally to LJ (34.1% vs 34.1; p=1.000) and MGIT (34.1% vs 50.7%; p=0.055). Notably, TLA had a lower incidence of uninterpretable results (4.9%) compared to LJ (7.0%) and MGIT (5.1%), although this difference was not stastically significant. Uninterpretable TLA results were primarily due to contamination (4 cases) or medium drying (3 cases), the latter attributed to the candle proximity during incubation. In contrast, LJ and MGIT encountered contamination as the primary issue.

**Figure 3.**
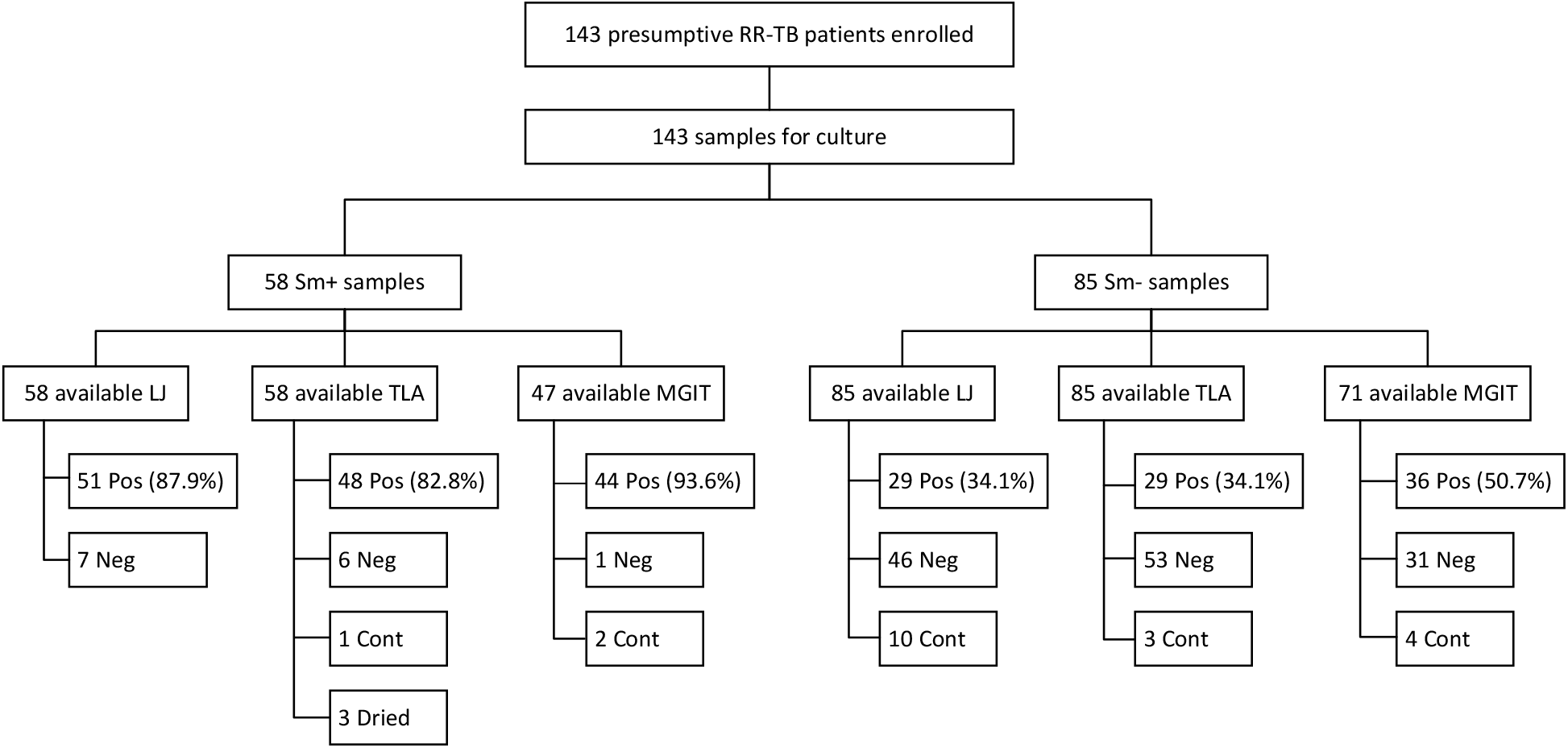
Sensitivity of direct-TLA-DST for primary MTB isolation by smear grade compared with other media. *RR-TB: rifampicin-resistant tuberculosis. Sm+: smear-microscopy-positive samples. Sm-: smear-microscopy-negative samples. LJ: Löwenstein-Jensen medium. TLA: thin-layer agar. MGIT: Mycobacteria Growth Indicator Tube. Pos: culture positive. Neg: culture negative. Cont: culture contaminated*.

For sm+ samples, TLA’s median time to MTB primary growth was 15 days (IQR 12-21), comparable to MGIT’s 14 days (IQR 10-23.5) and significantly faster than LJ’s 26 days (IQR 21-32). For sm-samples, TLA achieved MTB growth in a median time of 22 days (IQR 20-27), compared to 25.5 days for MGIT (IQR 19.5-32.75) and 44 days for LJ (IQR 30-55.5). Since TLA provided pDST results simultaneously with MTB isolation, it outperformed the other two culture-based pDST methods.

Among the conditions assessed in the TLA methodology, the incubation method did not exhibit a statistically significant effect on plates containing STC, regardless of smear status (p = 1 for sm+ samples and p = 0.28 for sm-samples) (Figure 4). In contrast, when comparing STC-containing plates to those without STC, the latter exhibited significantly better MTB isolation rates in sm− samples, achieving 71.4% versus 27% for STC plates (p = 0.0037). However, this distinction was less pronounced in smear-positive samples, where non-STC plates achieved 90.9% isolation rates compared to 80.9% for STC plates (p = 0.67). Further comparisons between the three experimental conditions, using LJ culture as a baseline, indicated that observed differences across conditions were unlikely to be due to random variation (p = 0.027 for sm+ samples and p < 0.001 for sm− samples).

**Figure 4.**
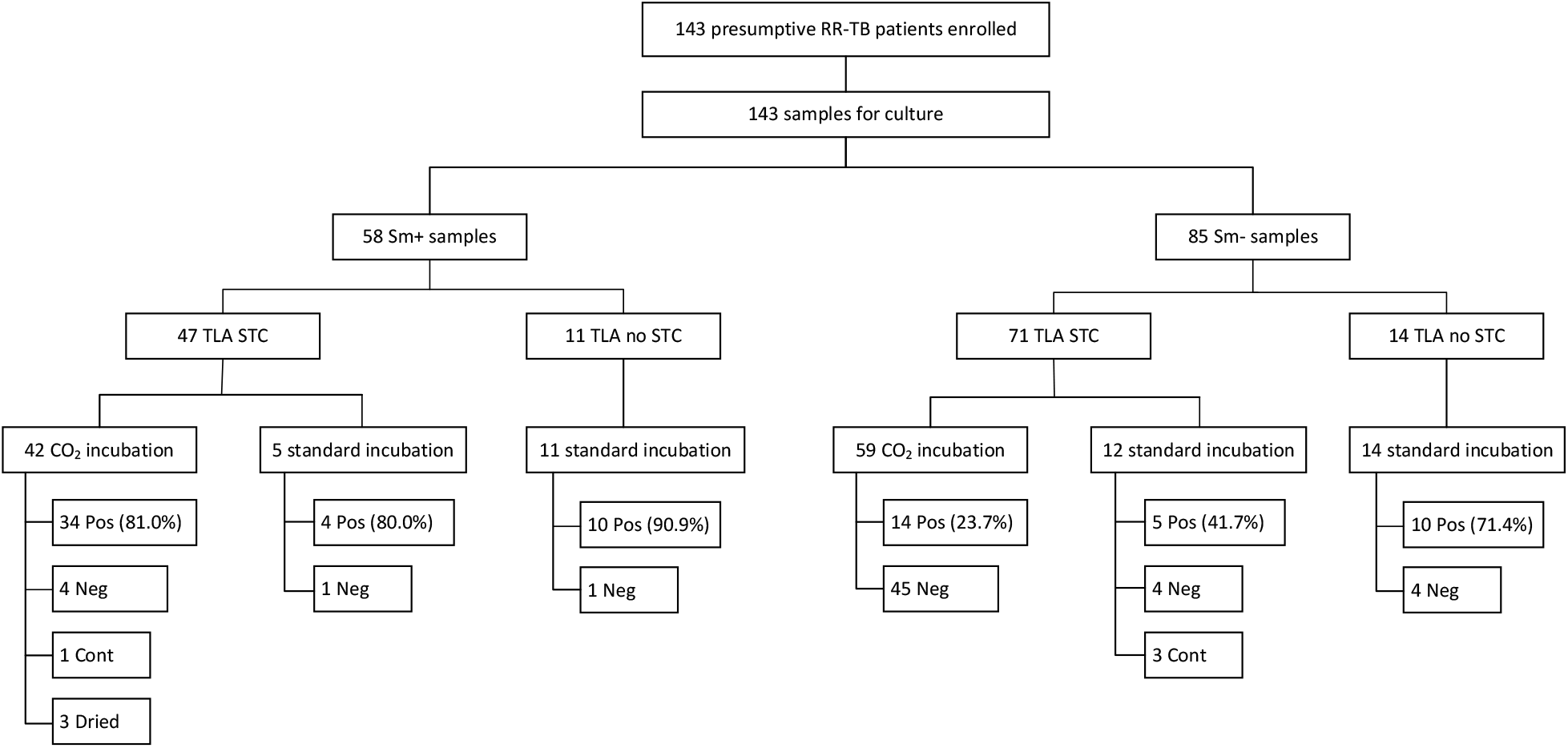
Flowchart showing enrolled patients and samples for direct-TLA-DST evaluation, stratified by smear-microscopy, STC addition and TLA plate incubation conditions. *RR-TB: rifampicin-resistant tuberculosis. Sm+: smear-microscopy-positive samples. Sm-: smear-microscopy-negative samples. TLA: thin-layer agar. STC:* 2,3-diphenyl-5-(2-thienyl) tetrazolium chloride. *Pos: culture positive. Neg: culture negative. Cont: culture contaminated. NI: non interpretable*

Of the 77 samples that showed growth in direct-TLA-DST, 67 had reference pDST results for comparison: 29 by LJ and 38 by MGIT. The specificity of TLA compared to indirect DST was 93.6% for RIF, 85.7% for INH, and 100% for both LFX and BDQ. The sensitivity was 95% for RIF and 84.6% for INH, while the sensitivity for LFX and BDQ could not be assessed due to the absence (BDQ) or an insufficient number (LFX) of resistant strains in the cohort. When discrepancies between direct-TLA and either indirect-MGIT or -LJ DST were examined against the reference standard, 13 out of 16 discordant results (81%) were resolved, with 9 (69%) favoring TLA (Table 2). The adjusted sensitivity for direct-TLA-DST was 100% for RIF and 87.8% for INH, while the specificity was 100% for all drugs, except INH (96.2%) (Table 3).

**Table 3.**
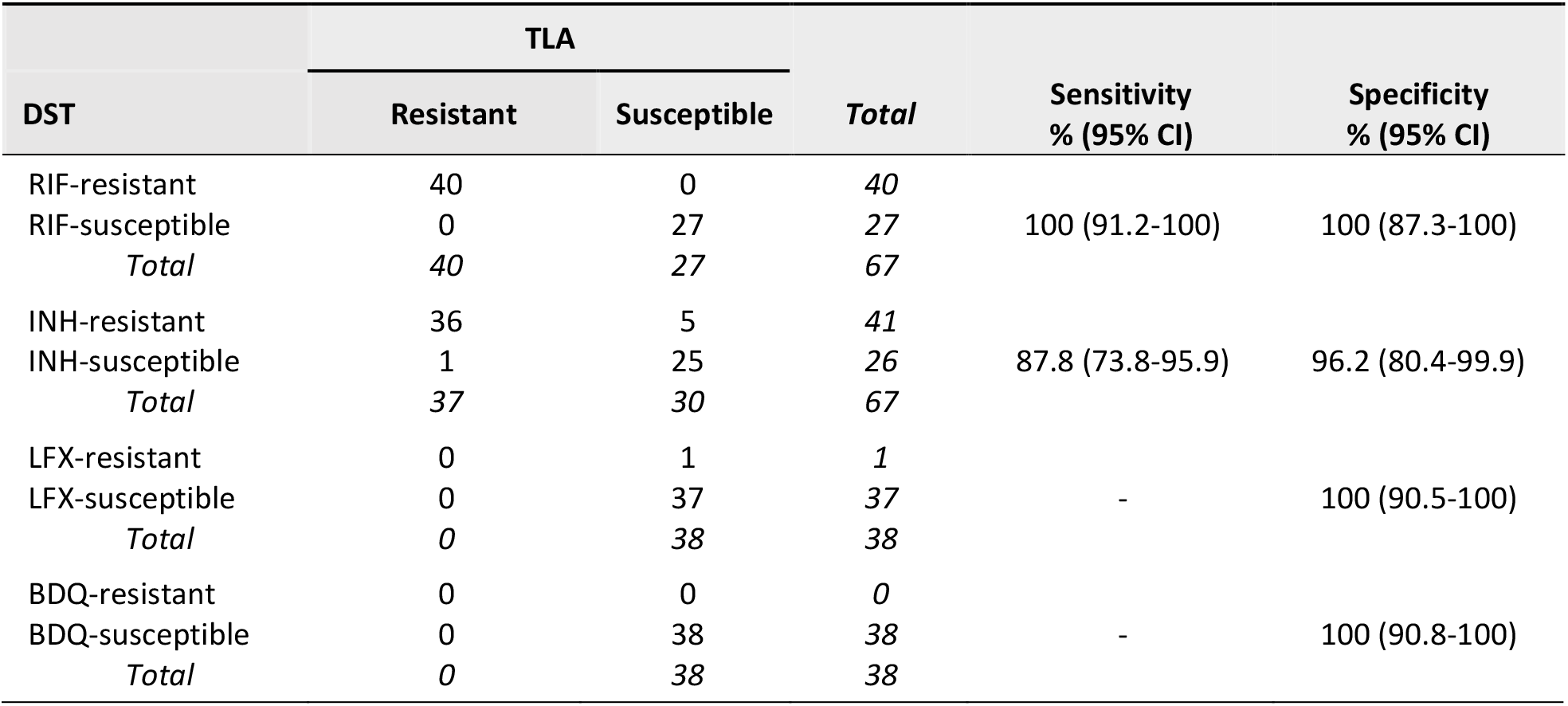
Comparison of drug-susceptibility results obtained by direct thin layer agar (TLA) testing, versus indirect testing on Löwenstein-Jensen (LJ) or in MGIT medium for rifampicin (RIF), isoniazid (INH), levofloxacin (LFX), and bedaquiline (BDQ) after resolving discrepancies. *DST: drug susceptibility testing. CI: confidence interval*.

## DISCUSSION

Based on our findings TLA performs well for indirect BDQ-MIC determination, as well as direct test for MTB isolation and DST for BDQ and other anti-TB drugs against established pDST methods on LJ and in MGIT. The BDQ-MIC values observed for the H37Rv strain by indirect-TLA confirmed its technical accuracy, with MICs consistently falling below the tentative CC of 0.25 ug/ml, and 99% inside the reference MIC quality control (QC) range for BDQ (0.015 to 0.12 ug/ml).^(16,17)^ For indirect testing, TLA based MIC determination saves time and consumables over tube based MIC determination (solid 7H11 or, by inference, liquid MGIT tubes), with reading at D7 showing optimal results. Moreover, the standard incubator outperformed the 5-10% CO_2_ incubator, further enhancing the feasibility of adopting TLA in routine diagnostic settings. While prior studies suggested that CO_2_ incubation might reduce the time to detection, our study found no significant difference in timing between the two methods, with results reliably available by day 7 in both conditions.^(24)^

TLA’s performance for MTB isolation and direct DST was further validated in this study. Consistent with previous research, TLA showed a high positivity rate for MTB detection in smear-positive samples, closely matching the 83% positivity rate reported by other studies.^(25,26)^ In smear-negative samples, TLA could isolate MTB comparably to LJ yet lower than MGIT, although the rate of uninterpretable results due to contamination was low. The low contamination rates and short detection times associated with TLA highlight its suitability for use in settings where rapid diagnosis – and potentially treatment follow-up - is essential for optimal patient management. Furthermore, transitioning from a CO_2_-enriched candle jar to standard incubation mitigated drying issues, further lowering the already minimal rates of uninterpretable results. This adjustment not only enhances TLA’s reliability but also its potential scalability to peripheral laboratories.

Direct-TLA-DST proved highly effective in detecting RIF resistance, showing robust sensitivity and specificity.^(19,20)^ However, the performance in detecting isoniazid (INH) resistance was somewhat lower than anticipated, with some unresolved discrepancies. Nonetheless, TLA’s ability to rapidly exclude INH resistance remains a key strength of the method.

This study is also the first to evaluate TLA as a direct method for detecting LFX and BDQ resistance. Although sensitivity could not be assessed due to an insufficient number of resistant samples, the high specificity observed for these drugs demonstrates TLA’s potential as a rapid diagnostic tool for ruling out extensively drug-resistant TB (XDR-TB) cases, representing a significant advancement for guiding MDR-TB treatment.

Our study presented some limitations. The relatively small sample size used to evaluate the TLA-BDQ-MIC should be expanded to include a broader range of MTB complex lineages, as lineages L5 to L10 were not represented. In the clinical setting, the stockout of reagents and the use of three different experimental conditions for direct-TLA-DST led to data stratification, which may have affected the interpretation of the results. Moreover, while TLA performed well in detecting resistance in susceptible cases, the limited number of resistant isolates for BDQ and LFX prevented a comprehensive assessment of TLA’s sensitivity to detect resistance to these drugs. Finally, some discrepancies between DST methods remain unresolved, highlighting the need for further investigations into the underlying causes of discordant results.

Overall, the results of this study contribute to the growing evidence supporting TLA as a valuable tool to rapidly diagnose resistance to bedaquiline in BSL2 level decentralized conditions. TLA’s ability to deliver accurate results within a short timeframe, combined with its simplicity, makes it an attractive option for rapid pDST of BDQ in laboratories with moderate biosafety requirements (level 2). Incorporating growth-enhancing components into the medium and/or increasing the inoculum volume could further improve TLA’s diagnostic efficacy, particularly for smear-negative samples.^(27,28)^ Additional research is required to validate TLA’s performance for BDQ sensitivity and extend its application to other new- and repurposed anti-TB drugs.

---

**Table S1.** Characteristics, genetic profile, and median BDQ-MIC values for described MTB isolates used in this study, comparing TLA and 7H11 methods.

| Isolate number | Lineage | Origin | *Rv0678* | *atpE* | Final confidence grading WHO | TLA MIC median | 7H11 MIC |
| --- | --- | --- | --- | --- | --- | --- | --- |
| 1999-01856 | Unknown | Clinical | Unknown | unknown |  | 0.25 | 0.25 |
| 2012-00594 | Unknown | Clinical | Unknown | unknown |  | 0.06 | 0.125 |
| 2014-01618 | Unknown | Clinical | Unknown | unknown |  | 1 | 1 |
| 2015-02105 | L4 | Clinical | WT | WT |  | 0.06 | 0.06 |
| 2015-02116 | L2 | Clinical | WT | WT |  | 0.03 | 0.03 |
| 2015-02119 | L2 | Clinical | WT | WT |  | 0.06 | 0.06 |
| 2015-02124 | L2 | Clinical | WT | WT |  | 0.06 | 0.03 |
| 2015-02137 | L2 | Clinical | WT | WT |  | 0.06 | 0.06 |
| 2015-02146 | L2 | Clinical | WT | WT |  | 0.03 | 0.03 |
| 2015-02147 | L2 | Clinical | p.Tyr157Cys | WT | Not present in catalogue 2<sup>nd</sup> ed | 0.25 | 0.25 |
| 2015-02150 | L2 | Clinical | WT | WT |  | 0.06 | 0.03 |
| 2015-02161 | L4 | Clinical | WT | WT |  | 0.06 | 0.03 |
| 2018-00082 | L1 | Clinical | WT | WT |  | 0.06 | 0.125 |
| 2018-00089 | L3 | Clinical | WT | WT |  | 0.06 | 0.06 |
| 2018-00090 | L3 | Clinical | WT | WT |  | 0.06 | 0.125 |
| 2018-00092 | L4 | Clinical | c.692C>T | WT | Not present in catalogue 2<sup>nd</sup> ed | 0.06 | 0.125 |
| 2018-00094 | L4 | Clinical | WT | WT |  | 0.06 | 0.03 |
| 2018-00102 | L1 | Clinical | WT | WT |  | 0.06 | 0.06 |
| 2019-00870 | L2 | Clinical | WT | WT |  | 0.06 | 0.06 |
| 2019-00875 | L2 | Clinical | p.Arg89Leu | WT | Uncertain significance | 0.5 | 0.5 |
| 2019-00918 | L2 | Clinical | WT | WT |  | 0.06 | 0.03 |
| 2013-00482 | L4 | In vitro selected | p.Arg135Trp | WT | Uncertain significance | 0.5 | 1 |
| 2013-02484 | L1 | In vitro selected | p.Tyr92STP | WT | Assoc w R - Interim | 1 | 1 |
| 2013-02485 | L1 | In vitro selected | p.Glu138Gly-<br>p.Met139Leu | WT | Not present in catalogue 2<sup>nd</sup> ed | 0.5 | 1 |
| 2014-01612 | L1 | In vitro selected | p.Ser63Arg | WT | Uncertain significance | 1 | 1 |
| 2014-01618 | L4 | In vitro selected | p.Arg135Trp | WT | Uncertain significance | 0.5 | 1 |
| 2014-02967 | L2 | In vitro selected | WT | p.Ala63Pro | Assoc w R - Interim | 1 | 1 |
| 2014-02973 | L2 | In vitro selected | WT | p.Glu61Asp | Assoc w R - Interim | 0.5 | 0.5 |
| 2014-02988 | L2 | In vitro selected | p.Gly78Val | WT | Not present in catalogue 2<sup>nd</sup> ed | 0.25 | 0.5 |
| 2014-03045 | L2 | In vitro selected | WT | p.Ala63Pro | Assoc w R - Interim | 2 | 2 |
| 2015-00554 | L4 | In vitro selected | WT | p.Asp28Gly | Assoc w R - Interim | 0.5 | 0.25 |

